# Plasticity of body axis polarity in *Hydra* regeneration under constraints

**DOI:** 10.1101/2021.02.04.429818

**Authors:** Anton Livshits, Liora Garion, Yonit Maroudas-Sacks, Lital Shani-Zerbib, Kinneret Keren, Erez Braun

## Abstract

One of the major events in animal morphogenesis is the emergence of a polar body axis. Here, we combine classic grafting techniques with live imaging to study the emergence of body axis polarity during whole body regeneration in *Hydra*. Composite tissues are made by fusing two rings, excised from separate animals, in different configurations that vary in the polarity and original positions of the rings along the body axes of the parent animals. Under frustrating initial configurations, body axis polarity that is otherwise stably inherited from the parent animal, can become labile and even be reversed. Importantly, the site of head regeneration exhibits a strong bias toward the edges of the tissue, even when this involves polarity reversal. In particular, we observe head formation at an originally aboral tissue edge, which is not compatible with models of *Hydra* regeneration based only on preexisting morphogen gradients or an injury response. Rather, we suggest that the structural bias toward head formation at the doublets’ edge is reinforced by the presence of a defect in the organization of the supra-cellular actin fibers, which invariably forms at the edge as the tissue heals. In this scenario, the defect supports head formation at the edge, even though a defect is neither required nor sufficient for head formation. Altogether, our results suggest that body axis determination is an integrated process that arises from the dynamic interplay between mechanical feedback and signaling processes.

## Introduction

The formation of a body axis is one of the fundamental initial steps of animal morphogenesis ^1^. The establishment of an axis leads the patterning process, delineating both the alignment and the polarity of the developing body plan. This is true both in developing embryos and during whole body regeneration in organisms such as planarians or *Hydra*. Whereas in some cases the establishment of a body axis occurs *de novo* and involves a spontaneous symmetry breaking event ^2, 3^, often there is a strong memory of polarity inherited from the parent animal. The memory of polarity is evident in excised tissues of planaria ^4^ or *Hydra* ^5, 6^, which typically regenerate along the direction of the original body axis, as well as in developing animals such as drosophila where polarity arises from inherited gradients of maternal mRNA in the oocyte ^1^. Interestingly, the molecular toolbox underlying axis formation in embryonic development and during regeneration is largely shared among different animals, with a central role for the Wnt signaling pathway ^1, 3, 4, 7–10^. Despite extensive research over the last few decades, many fundamental questions regarding body axis formation and maintenance in animal morphogenesis remain largely open.

Here we focus on body axis polarity in the small fresh-water animal *Hydra*. *Hydra* have a single polar body axis, with a head at one end and a foot at the other end. This simple uniaxial body plan together with *Hydra*’s remarkable regeneration capabilities make it an excellent model system to explore fundamental concepts related to the establishment and maintenance of body axis polarity. Indeed, classic work in *Hydra* was instrumental in shaping our current understanding of axial patterning, establishing the important role of an “organizer” ^6, 11^, and putting forward the idea of pattern formation by reaction-diffusion dynamics of morphogens^12–14^ and the concept of positional information ^15–17^.

Extensive previous work has shown that the body axis – both its alignment and its polarity – are strongly preserved in regenerating *Hydra*. The memory of polarity in *Hydra* was first demonstrated by Abraham Trembley, who in 1744 showed that bisected *Hydra* can regenerate a head or foot according to their original polarity ^18^. Later work showed that the memory of polarity is retained even in small excised tissue segments ^5, 19^. Despite this memory, the polarity of regenerating *Hydra* tissues can also exhibit substantial plasticity under certain conditions ^20–23^. For example, gastric tissue pieces can exhibit polarity reversal in response to local pharmacological perturbations ^23^ or be induced to switch polarity by grafting a head and a foot from another animal in the opposite polarity ^21, 22^.

A central player in the maintenance of the polar body axis in *Hydra* is the head organizer, located at the tip of the hypostome in the head of mature animals ^6, 11^. The organizer activity is thought to be autocatalytic and produces both activation and inhibition signals that form morphogenetic gradients originating from the head and extending throughout the animal’s body. The activation signal promotes the formation of a new head whereas the inhibitory signal represses the formation of a second head near an existing head, and restricts bud formation to the lower part of the body, closer to the foot side. The position-dependent activation and inhibition activity and their propagation throughout the *Hydra* body were extensively investigated in a series of grafting experiments in which the influence of the grafted tissue was studied as a function of the original position of the tissue in the donor animal and the grafting position in the host animal (reviewed in ^24^).

During regeneration from tissue segments, following the removal of the head and foot regions, a new head organizer has to emerge. While the entire gastric tissue has the potential to form a new organizer, the realization of this potential typically occurs only in the most oral part of the excised tissue ^5^. The establishment of a new organizer is thought to occur via a local self-enhancing reaction that depends primarily on a preexisting gradient of activation properties of the tissue (sometimes termed “competence”) ^14, 25^. While the molecular basis for this graded head activation potential is still not entirely clear, work over the last few decades has elucidated many of the molecular factors associated with organizer formation and polarity determination in *Hydra* ^3, 7, 8, 10, 26, 27^. In particular, the most prominent pathway associated with the head activator is the Wnt pathway, with the expression of Wnt3 at the tip of the hypostome appearing to be one of the earliest indications for the formation of a new head organizer ^3, 8, 26, 28^. Perturbations that globally upregulate the Wnt pathway, for example by overexpressing a β-catenin transgene ^29^ or by pharmacological stabilization of β-catenin by alsterpaullone ^30^, have been shown to induce multiple head organizers across the entire *Hydra* body column.

The injury generated during excision of *Hydra* tissues has also been shown to have an important role in head regeneration. Injury has been shown to promote head regeneration in regeneration-deficient *Hydra* strains ^31, 32^, whereas the lack of injury in animals that were carefully bisected without wounding by ligature interfered with head regeneration ^33^. More recent experiments at the molecular level revealed the extensive signaling events triggered by injury ^7, 10, 26, 27, 34^. In particular, the Wnt pathway was found to be activated at the wound site following injury ^10, 26, 27, 34^. The link between the rapid activation of signaling pathways following injury and the subsequent patterning that eventually leads to regeneration is however still unclear, and in particular it is not known what discriminates between the oral and aboral sides of the wounded tissue ^7, 27, 35^. An earlier study reported a position-dependent Wnt3 enhancement near the oral wound site via an apoptosis-mediated mechanism ^34^, yet several recent experiments failed to find early signatures of a position-dependent injury response, detecting activation of apoptosis and enhanced biosignaling in both the oral and the aboral wound sites ^10, 27, 35^.

In addition to polarity information associated with preexisting tissue gradients and the injury response, there is also some evidence suggesting that structural factors can play a role in the formation of a new head organizer. Early work indicated that the local structure ^36^ and the shape of the closure region of the injury site ^5^ can affect the potential for head formation. More recently, we found that the cytoskeletal organization plays a role in the establishment of the body axis during regeneration from excised tissue pieces ^37, 38^. We showed that the parallel array of ectodermal supra-cellular actin fibers in the parent animal remains partially intact in excised tissue segments, providing a structural memory of the alignment of the body axis in the parent *Hydra* that defines the alignment of the body axis in the regenerating animal ^37^. In subsequent work, we showed that local defects in the alignment of these actin fibers act as organization centers for morphogenesis, with morphological features appearing at defect sites ^38^. In particular, we showed that head formation in regenerating tissue fragments occurred at the site of an aster-like defect in the alignment of the ectodermal actin fibers.

Despite this progress, the mechanisms involved in the memory of polarity and the dynamics leading to the establishment of a new organizer in regenerating *Hydra* are still unclear. Here we combine classic experimental design with modern imaging techniques to gain insight into this important process by exploring the flexibility and reorganization capacity of polarity under constraining conditions. We study the regeneration of composite tissues generated by careful adhesion of two excised tissue rings in configurations in which the rings’ polarities are not mutually aligned and/or their relative positions along the parent animals’ body axis are rearranged. As such, our experiments are designed to introduce frustration in the biochemical and structural factors associated with body axis polarity, so the polarity of the fused ring doublets cannot be simply defined by the original polarity of the excised tissues. We follow the regeneration dynamics and cytoskeletal organization of the composite ring doublets by live microscopy. The regenerating doublets display higher flexibility to reorganize their body axis compared to bisected animals (that retain either a head or a foot and maintain their original polarity), allowing us to expose the dynamic competition between different processes involved in body axis determination in *Hydra*.

We find that under certain frustrating configurations, the original tissue polarity can be reversed, with a head regenerating from the aboral edge of one of the excised rings. Such polarity reversal demonstrates the plasticity of the polarity establishment process and is not compatible with models assuming that body axis polarity is determined by a single causal factor; be it preexisting morphogenetic gradients in the excised tissue, wound healing at an oral injury site, or another inherited factor from the parent animal. Rather, our experiments reveal the ability of tissue polarity to rearrange dynamically after resetting its previous memory to accommodate the imposed constraints. Our results further suggest that mechanical feedback and cytoskeletal organization can also contribute to the establishment of polarity in regenerating Hydra, aligned with our recent observations ^37, 38^. Overall our experiments complement the large body of classical work on polarity in *Hydra*, and allow us to offer a dynamic view of polarity determination during regeneration, which involves the integration of multiple factors including signaling processes, either inherited (based on the original polarity and position of the excised tissue) or emerging, and mechanical processes that depend on cytoskeletal organization and the tissue architecture^39^.

## Results

### Regeneration under frustrating conditions depends on tissue structure and polarity memory

Excised *Hydra* tissue rings retain a strong memory of their original polarity, regenerating a head on their original head-facing side and a foot on their original foot-facing side ^5^. We have confirmed these previous observations by studying excised *Hydra* tissue rings which had their oral side labeled, or alternatively by placing regenerating tissue rings on a thin wire ^37^ that marks their original orientation (Fig. S1). In both cases, we found that the original tissue polarity is maintained (Fig. S1). As in bisected animals, the polarity memory in a regenerating ring is inherited in a straightforward manner-the most oral region in the original tissue develops into the head of the regenerated animal, whereas the aboral side at the opposite edge becomes the foot.

To explore the interplay between polarity memory and structure we set out to generate frustrating initial configurations in which the tissue polarity cannot be maintained in a simple way. This is achieved by fusing two excised rings (originating from two separate animals) along their axis to form a single composite tissue. The configuration of the composite ring doublet can be modulated in a controlled manner by varying the orientation of the two fused rings relative to their original polarity and/or by varying their position relative to their original location along the parents’ body axes. To generate the composite tissue, the excised rings are threaded in tandem on a wire ^37, 40^ in the desired configuration, maintained there for ∼6 hours to allow the tissues to adhere, and subsequently removed from the wire and followed by time lapse microscopy (Methods). To enable tracking of the tissue dynamics, the two rings are excised from separate animals that are differentially marked by electroporating distinct fluorescent cytosolic dyes (Methods).

In the Head-to-Head (H2H) configuration, two rings are placed opposing each other so the originally head-facing sides of the two rings adhere (Fig. 1A). In this configuration, the memory of the original polarity favors head formation in the middle, whereas the structure of the fused ring doublet resembles that of a (single) thicker ring in which the open ends seal and a head regenerates at the oral edge. The majority of fused ring doublets in the H2H configuration regenerate and form a head (i.e. a hypostome with two or more tentacles) (145 out of 164 samples). The outcome morphologies of the regenerated animals are characterized using the fluorescent labeling of the ring tissues to identify the site of new head formation (Figs. 1B, S2). The majority (85/145) of regenerated ring doublets formed a head in the middle, as expected based on the original tissue polarity (Figs. 1C, S2ii,iii; Movie 1). In these cases, the new organizer forms at the interface between the two rings as indicated by the fluorescent labels which are split across the regenerated head (Figs. 1C, S2ii,iii). The two edges of the ring doublet typically turned into two feet (Figs. 1C, S2ii), but in some cases the two ends merged to form a single foot (Fig. S2iii).

**Fig 1.**
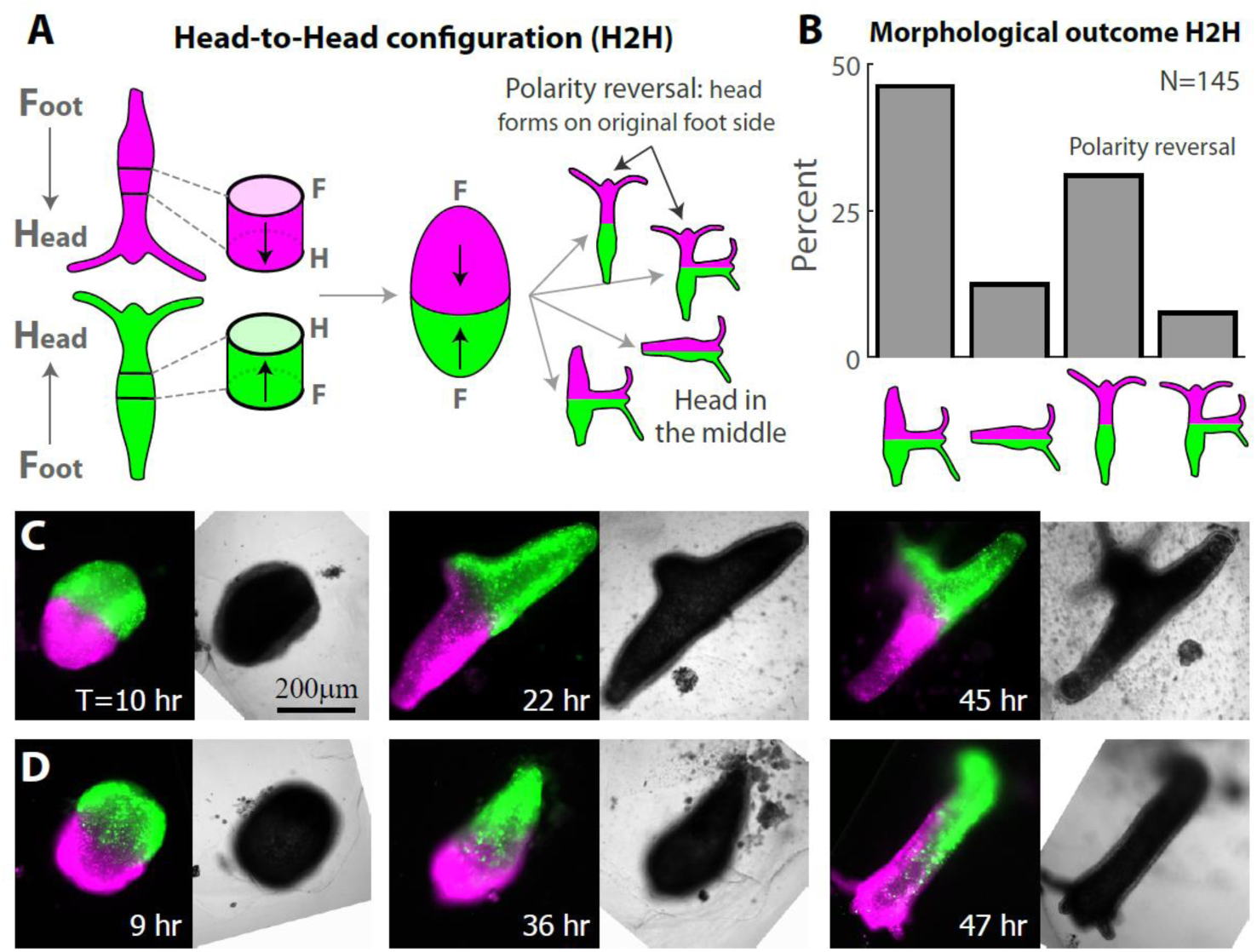
Polarity rearrangement in regenerating Head-to-Head (H2H) ring doublets. (A) Schematic illustration of the regeneration experiments with fused ring doublets in the H2H configuration. Two rings are excised from different animals (labeled in green or magenta). The rings adhere to each other at their head-facing sides, and the regeneration of the fused ring doublet is followed over time. (B) Bar plot depicting the outcome morphologies of fused H2H ring doublets. Experiments were performed on 164 samples. 2/164 disintegrated, 17/164 did not regenerate, and 145/164 regenerated. The outcome morphologies of the regenerated animals were categorized as: a head in the middle with two feet, a head in the middle with one foot, a head at either edge (green or magenta), a head in the middle and at the edge (Methods; Fig. S2). All but 4/145 (i.e. 97%) of the regenerated animals fell into one of these categories. The percentages of regenerated animals in each category are depicted. (C) Images from a time lapse movie of a H2H doublet that regenerated into an animal with a head in the middle (Movie 1). (D) Images from a time lapse movie of a H2H doublet that underwent polarity reversal, regenerating into a normal morphology with a head developing from an originally foot-facing side of one of the excised rings (Movie 2). The panels in (C,D) depict combined epifluorescence (left; green-AlexaFluor 647-conjugated 10kD dextran, magenta-Texas Red-conjugated 3kD dextran) and bright-field (right) images at the specified time points during the regeneration (time after excision in hours).

Surprisingly, a considearble fraction of regenerating *Hydra* in the H2H configuration formed a head at the edge of the ring doublet, i.e. at the region that was farthest from the head in the parent animals (Fig. 1D; Movie 2). This was observed in more than 30% of the samples (45/145), where a head formed at an aboral wound site on an originally foot-facing side of one of the excised rings. Occasionally, the regenerating animals formed both a head in the middle and at the edge (11/145; Figs. 1B, S2v). The outcome morphologies, and in particular the propensity for new head formation at the middle versus the edge of the ring doublet, did not exhibit a correlation with the relative sizes of the two tissue rings (Fig. S3).

The lack of head formation at the middle in a considerable fraction of the H2H samples (Fig. 1B,D), suggests that the fusion of the second ring at the oral wound site has a repressing effect on head formation. To examine the influence of the interface between the two rings on head formation, we designed experiments in which we purposefully added a physical perturbation at the adhesion interface by placing a 75 μm-diameter wire at the fusion interface for ∼6 hours (Fig. S4C; Methods). The transient introduction of this perturbation at the interface increased the fraction of samples that preserved their polarity and regenerated a head in the middle (86% = 24/28 of the samples, compared to 66%=96/145 without a wire; Fig. S4D). The increase in the fraction of ring doublets that form a head at the interface following this transient perturbation is aligned with earlier observations by Shimizu and Sawada that showed that structural abnormalities at the graft site correlated with additional head formation in tissue transplantation experiments in mature *Hydra* ^36^.

### The influence of the original tissue position on polarity determination

The potential to regenerate a head and form a new organizer is present in the entire gastric tissue in *Hydra*. Yet, the head activation potential appears to be graded along the body axis, decaying away from the head organizer ^25^. This was illustrated by classic grafting experiments which demonstrate that the probability to form a new head increases when the source of the grafted tissue is closer to the head of the donor animal ^24, 41^. The graded head activation potential is also reflected in the time it takes to regenerate a head, which increases if the site of decapitation is further down the body column ^19, 42^. The position-dependent regeneration potential of *Hydra* tissues was further demonstrated by experiments by Sugiyama and coworkers who formed elongated cylindrical tubes by fusing a large number of tissue rings, and showed that the potential to form head structures depended on the original location of the excised rings along the body axis of the parent animal ^40^. These observations motivated us to utilize the original position of the excised tissue along the axis of the parent animal as an additional control parameter and ask: What are the implications of the graded head activation potential in the parent animals on the plasticity and reorganization dynamics of polarity in regenerating tissues under frustrating conditions?

The regeneration outcome in the H2H configuration reflects a competition between the memory of the original tissue polarity, and the constraints imposed by the fusion of the two rings to each other. This competition masks the influence of the position of the excised tissue in the parent animals (Fig. S4). To focus on the effect of the original tissue position, we turned to ring doublets in the Head-to-Foot (H2F) configuration, placing the two excised rings in the same orientation by adhering the oral end of one ring to the aboral side of the second ring, while varying the original positions of the rings in the parent animals (Fig. 2A). The rings are excised from different tissue locations by bisecting a *Hydra* in the middle and taking either the **U**pper ring (above the midline; **U**) or the **L**ower ring (below the midline; **L**) (Fig. 2A; Methods). The ring doublets are then fused in the H2F configuration, in an *oriented* manner that maintains their relative positions along the body axis (Fig. 2A top), or in an *anti-oriented* manner with the upper ring positioned below the lower ring (Fig. 2A bottom). Note that in both configurations, the first ring faces an additional ring on its oral interface, while the second ring has a free edge on its oral side. Importantly, unlike the H2H configuration in which the two rings have opposite initial polarities, in the H2F configuration the fused doublet has a preferred orientation along which it can regenerate while maintaining the original polarity of both rings.

**Figure 2.**
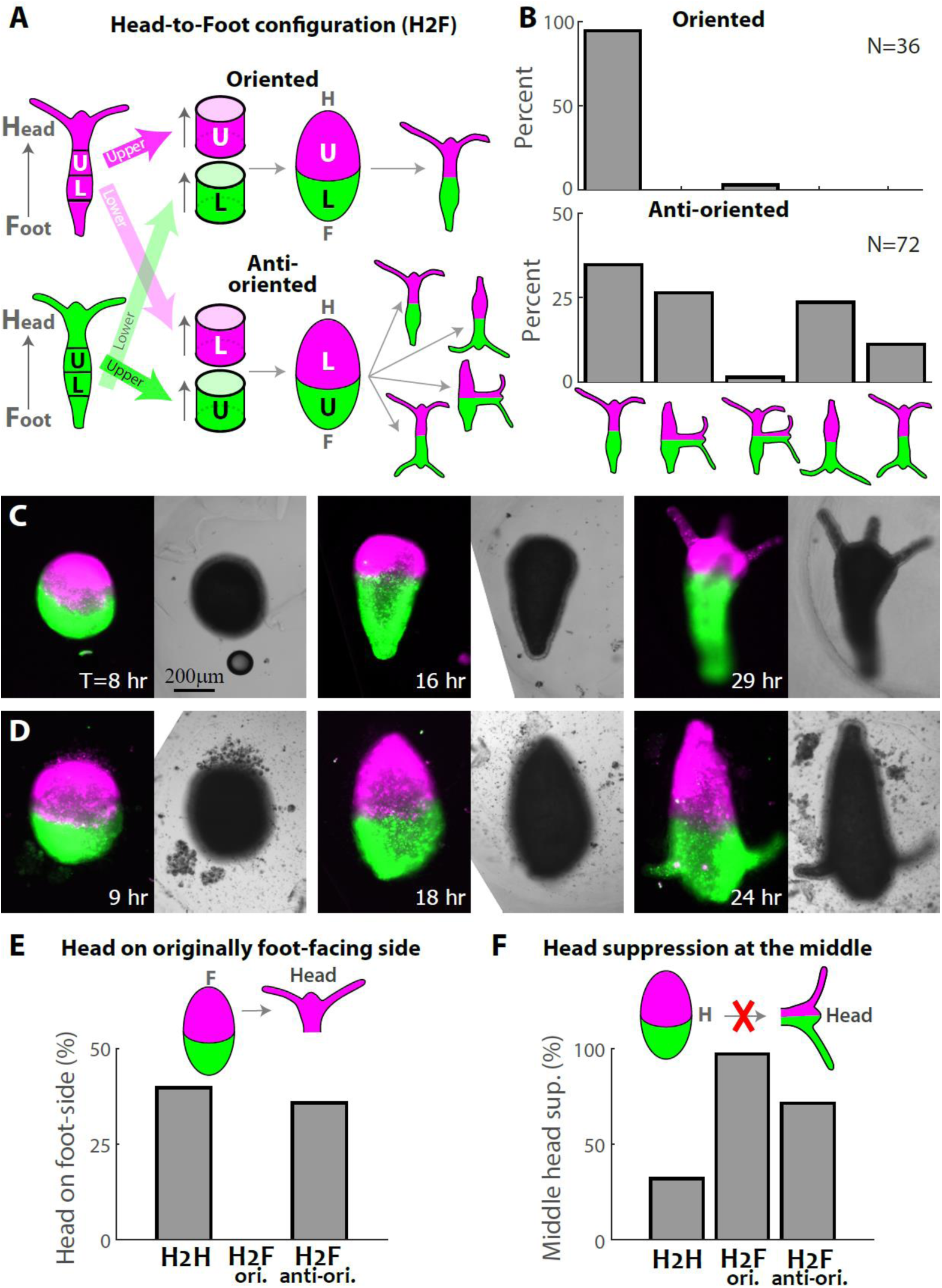
Regeneration of Head-to-Foot (H2F) ring doublets depends on the position of the excised rings within the parent animals. (A) Schematic illustration of the regeneration experiments with fused ring doublets in the H2F configuration. Rings are excised from above (**U**) or below (**L**) the approximated midpoint of the two parent animals that are differentially labeled (green/magenta). The rings are fused so that the head-facing side of the first ring adheres to the foot-facing side of the second ring in an oriented configuration that preserves the relative position of the rings (top) or an anti-oriented manner (bottom). (B) Bar plot depicting the outcome morphologies of fused ring doublets generated in the H2F configuration in an oriented (top) or anti-oriented manner (bottom). Experiments were performed on 37 oriented ring doublet samples and 78 anti-oriented samples. In the oriented samples 1/37 disintegrated and 36/37 regenerated, out of which 34 regenerating along the original polarity. In the anti-oriented no samples disintegrated, 6/78 did not regenerate, and 72/78 regenerated into one of several possible outcome morphologies: a normal morphology along the original polarity, a head in the middle, a head in the middle and at the oral edge, a normal morphology with *flipped* polarity, two heads at the oral and aboral edges (Methods). All but 2/78 (i.e. 97%) of the regenerated animals fell into one of these categories. The percentages of regenerated animals in each category are depicted. (C) Images from a time lapse movie of an oriented H2F doublet that regenerated into a normal animal along the original body axis orientation (Movie 3). (D) Images from a time lapse movie of an anti-oriented H2F doublet that regenerated into an animal whose body axis is oriented in the opposite direction to its original polarity, with a head forming on the original foot-facing side (Movie 4). The panels in (C,D) depict combined epifluorescence (left; green-AlexaFluor 647-conjugated 10kD dextran, magenta-Texas Red-conjugated 3kD dextran) and bright-field (right) images at the specified time points during the regeneration. (E) Bar plot summarizing the incidence of head formation on an originally foot-facing edge (polarity reversal) in regenerating ring doublets in the H2H and H2F oriented and anti-oriented configurations. (F) Bar plot summarizing the incidence of head suppression at an originally oral ring interface in the middle of regenerating ring doublets in the different configurations. The data in (E, F) is based on Figures 1B,2B.

The regeneration of H2F ring doublets was found to dramatically depend on the relative tissue positions of the excised rings in the parent animals (Fig. 2). The regeneration of ring doublets that were fused in an oriented manner, preserving both the polarity and the relative position of the excised rings along the body axis, invariably resulted in normal animals, with a head regenerating on the original head-facing side (the U side) at one end of the doublet and a foot forming on the original foot-facing side at the other edge (the L side; Fig. 2B top, 2C; Movie 3). Note that none of the samples developed a head in the middle, despite the presence of an oral wound there, illustrating again (as in the H2H configuration) the repressing effect of additional tissue at an originally oral-facing interface (Fig. 2F).

Ring doublets that were fused in an anti-oriented manner displayed markedly different behavior, with variable morphological outcomes and a high incidence of polarity reversal (Fig. 2B bottom). Only about a third of the regenerated samples (25/72) developed a normal morphology along the direction of the original body axis, whereas a considerable fraction (17/72) of samples formed a head on the foot-facing side of the upper ring, thus regenerating into an animal with a polarity that is *flipped* relative to the original body axis of both rings (Fig. 2D; Movie 4). In both of these cases, the regenerated animal has a normal morphology, either along the original polarity of the excised rings or in the opposite orientation. In other cases, as in the H2H configuration, the frustration caused by the presence of the second ring and the competition between different sites, is resolved by alternative configurations with a head in the middle (19/72) or two heads (9/72; Fig. 2D).

These results highlight the influence of the original tissue position along the body axis of the parent animal as an important factor, in addition to the original polarity and structure of the regenerating tissue, in determining the polarity of the regenerated animal. The naively expected regeneration trajectory in the H2F configuration is the formation of an animal that preserves the original polarity in both rings, developing a head at the free oral edge. This is indeed the case in the oriented H2F configuration, where the original polarity is maintained in nearly all the samples. In contrast, the majority of the anti-oriented H2F samples do not follow this trajectory, revealing the importance of the original position of the tissue along the body axis of the parent animal in the processes involved in polarity determination. In some cases, as in the H2H configuration, the structural repression of a head in the middle is not strong enough and the frustration is resolved by forming a head in the middle (as a single head or in addition to a head at the edge). The truly unexpected result is the anti-oriented samples which regenerate into a normal morphology with a flipped polarity (Fig. 2D); despite having a free oral edge, these samples regenerate by growing a head on the opposite side, on an originally-aboral edge. This exposes a dynamic aspect of the establishment of polarity that was not appreciated before, involving an interplay between position-dependent regeneration kinetics and structural factors that jointly lead to this surprising outcome in a large fraction of the samples.

### Position-dependent inhibition of second head formation

The formation of two heads in regenerating fused ring doublets is infrequent, even though separately the two excised rings would each regenerate a head on their oral side. The suppression of the formation of a second head occurs even in the H2F configuration, in which the composite tissue contains two distinct regions with an oral interface (Fig. 2B), suggesting that the formation of one head in a regenerating tissue inhibits the formation of the second head ^43^. To examine the inhibition of second head formation during regeneration and the factors influencing the choice of the head-forming site, we turned to ring doublets in the Foot-to-Foot configuration (F2F; Fig. 3). In this configuration, the two rings adhere on their aboral ends, so the composite tissue has two free oral ends at both sides, allowing in principle the formation of two heads at both edges of the ring doublet.

**Figure 3.**
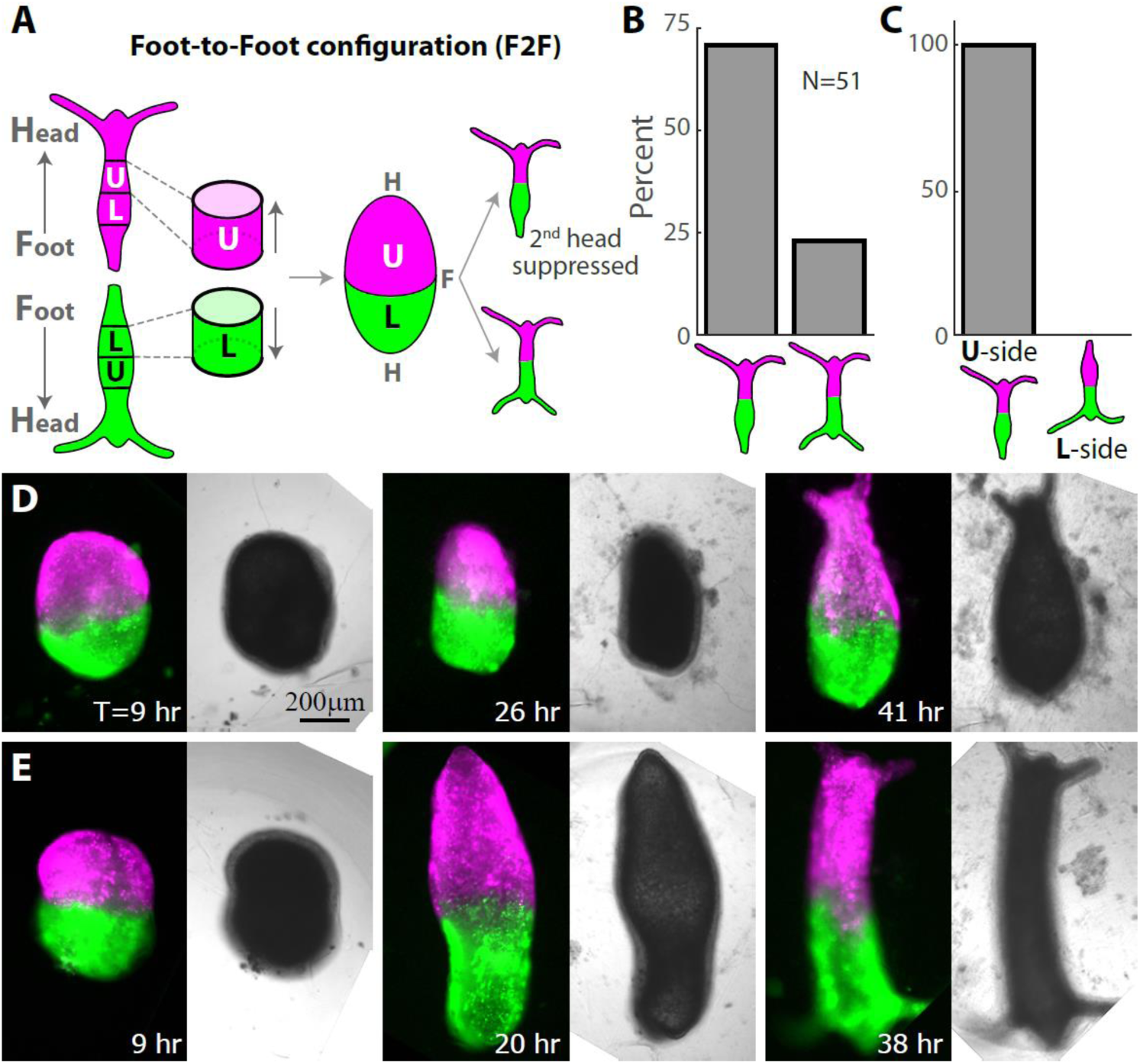
Position-dependent inhibition of second head formation in regenerating Foot-to-Foot (F2F) ring doublets. (A) Schematic illustration of the regeneration experiments with fused ring doublets in the F2F configuration. Rings are excised from above (U) or below (L) the approximated midpoint of animals that are differentially labeled (green/magenta). Two rings from different positions (U and L) are fused to each other at their foot-facing sides, and the regeneration is followed over time. (B) Bar plot depicting the outcome morphologies of fused ring doublets generated in the F2F configuration. Experiments were performed on 53 samples. 1/ 53 disintegrated, 1/53 did not regenerate and 51/53 regenerated into either a normal morphology with a single head, or an animal with two heads at both edges. All but 2/51 (i.e. 96%) of the regenerated animals fell into one of these categories. The percentages of regenerated animals in each category are depicted. (C) Bar plot showing the bias in head formation depending on the original position of the excised tissue rings (U or L). The original position of the excised ring on the side of the tissue where a head appeared was recorded for all F2F ring doublets that regenerated into an animal with a single head. In all these samples (36/36), invariably, the head formed on the U side. (D) Images from a time lapse movie of a F2F ring doublet that regenerated into an animal with a single head on the U side (Movie 5). (E) Images from a time lapse movie of a F2F ring doublet that regenerated into an animal with two heads on both sides (Movie 6). The panels in (D,E) depict combined epifluorescence (left; green-Texas Red-conjugated 3kD dextran, magenta-AlexaFluor 647-conjugated 10kD dextran) and bright-field (right) images at the specified time points during the regeneration (time after excision in hours).

We find that the majority of F2F doublets developed into animals that have a normal morphology with a head on one side and a foot on the other side (37 out of 51 samples that regenerated; Fig. 3B,D; Movie 5), with a minority regenerating two heads on both sides (12/51; Fig. 3E; Movie 6). Thus, even though there is no structural barrier for head formation at both ends of the composite tissue, the formation of a second head is frequently suppressed. This inhibitory effect is consistent with previous observations of the inhibition of head formation in the vicinity of clusters of head-activating cells in regenerating aggregates ^43, 44^.

To examine the position-dependent factors associated with this suppression we focused on asymmetric F2F ring doublets, combining rings from the regions above and below the midpoint (Fig. 3A). We find that head formation always occurs on the side that was originally closer to the head in the parent animals (the **U**-side; Fig. 3C). We do not observe foot formation in the middle, indicating that the presence of a second ring represses foot development, in analogy to the suppression of head formation in the middle in the H2H and H2F configurations (Fig. 2F).

These results illustrate again the influence of the original tissue position along the body axis of the parent animal on head formation; of the two original oral wound sites at the edges of the fused ring doublet, the site that was originally closer to the head “wins” (i.e. regenerates a new head), whereas head formation at the second site is typically suppressed (Fig. 3B,C).

### Cytoskeletal reorganization during doublet regeneration and the role of defects

The outcome morphologies of regenerating ring doublets in the different configurations highlight the contribution of structural factors to polarity determination during regeneration. In particular, we find a bias that favors head formation at the edge of regenerating ring doublets (Fig. 2E) and tends to suppress head formation in the middle (Fig. 2F). An important feature that distinguishes the edges of fused ring doublets from the rest of the tissue is the local cytoskeletal organization. Mature *Hydra* contain ectodermal actin myonemes that form a parallel array of supra-cellular fibers that span the entire length of the animal ^37, 45^. The myonemes are aligned with the animal’s body axis but are not expected to be polar, since these contractile fibers bundle together multiple actin filaments with opposite polarities. The organization of the supra-cellular actin fibers in a mature *Hydra* contains a characteristic set of defects, which are local regions in which the fibers are not mutually aligned. Importantly, the sites of these defects coincides with the morphological features in the animal ^38^. In particular, the organizer site at the tip of the mouth and the foot of the animal are characterized by aster-like defects in the alignment of the actin myonemes.

An excised tissue ring inherits the parallel myoneme array from the parent animal ^37^. Single regenerating rings typically seal the top and bottom edges of the ring (unless they undergo buckling ^37, 46^), to form a closed spheroid. As a tissue ring seals at its ends, the actin fibers aligned parallel to the ring’s axis converge at both ends into a small region with a characteristic aster-like organization of fibers at the edges (also known as a +1 nematic topological defect; Fig. 4). As shown below, the edges of composite ring doublets seal in a similar manner and therefore also contain an aster-like arrangement of actin fibers. Since polarity in individual regenerating rings is preserved, the regions around the aster-like defects ^38^ at the edges of the originally head or foot facing-side of an excised ring, should develop, respectively, into the head or foot of the regenerated animal (Fig. 4A). To directly demonstrate this, we follow the actin organization in regenerating tissue rings expressing lifeact-GFP that labels the actin filaments in the ectoderm layer ^37, 38, 45^ and mark the oral side of the excised tissue by locally uncaging a fluorescent dye there ^38^ (see Methods). Indeed, we find that the edges of a sealed ring have an aster-like defect (that can be localized at a point, or spread out over a small region) and that the tissue polarity is preserved with a head regenerating in the labeled region at the originally oral side of the ring (Fig. 4B; Movie 7). In particular, the region surrounding the defect at the oral wound becomes the site of the new head organizer at the tip of the mouth of the regenerated *Hydra* (Fig. 4B). Since the location of the aster-like defect coincides with the signaling center at the oral wound, it is difficult to assess the contribution of the presence of the defect for head organizer formation in this case.

**Figure 4.**
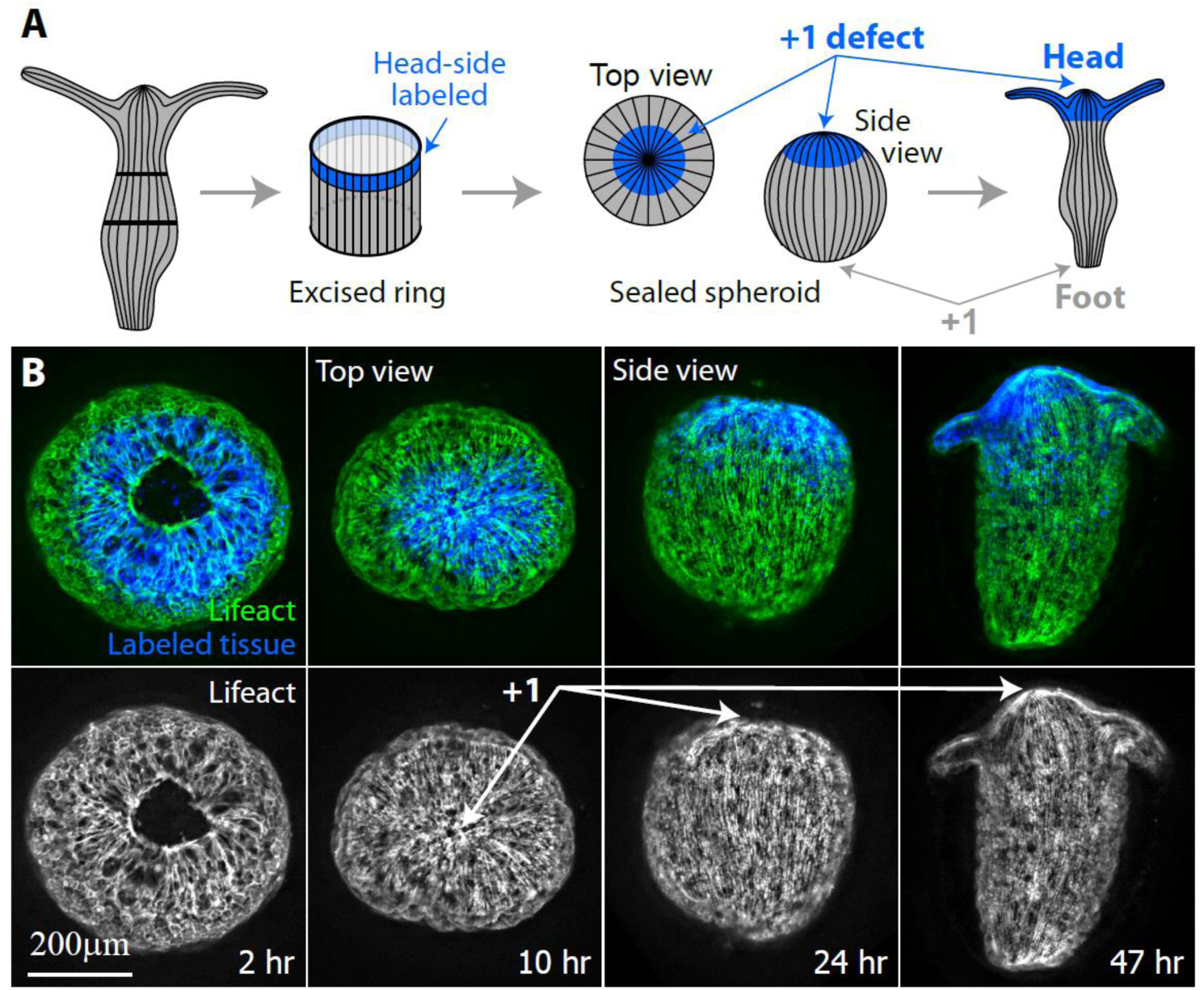
Actin fiber organization in a single regenerating ring. (A) Schematic illustration of the ectodermal actin fiber organization during regeneration of an excised ring. Left: The actin fibers in a mature *Hydra* are arranged in a parallel array, along the direction of the body axis. Middle left: An excised ring inherits this actin fiber organization. The region at the apical end of the excised ring is labeled (blue). Middle right: the excised ring closes into a spheroid by sealing the open ends at its top and bottom sides. Two aster-like defects form at both ends of the sealed spheroid. Right: The excised ring regenerates into an animal with a new head forming at the defect site located at the originally head-facing (labeled) side of the excised ring, and a new foot forming at the other end. (B) Spinning-disk confocal images from a time-lapse movie of an excised ring that is labeled at its apical end (Movie 7). The labeling is done by locally uncaging an electroporated caged-dye (Abberior CAGE 552; see Methods) (32). Images are shown at frames providing a top view of the regenerating ring from its apical end (left images) or a side view of the tissue (right images) at different time points during the regeneration process. Bottom: The projected lifeact-GFP signal (Methods) showing the organization of the ectodermal actin fibers. The aster-like (+1) defect at the apical side of the sealed spheroid is clearly visible from the top view, and coincides with the site of formation of the new organizer at the tip of the hypostome of the regenerated animal. Top: Overlay of the lifeact-GFP signal (green) and the fluorescent tissue label (blue) that marks the tissue originating from the apical side of the excised ring.

How do the actin fibers organize during regeneration of fused ring doublets? We follow the dynamic organization of the ectodermal actin fibers in regenerating doublets expressing lifeact-GFP ^37, 38, 45^, together with fluorescent probes labeling the tissue originating from either of the fused rings in the doublet (Methods). Following fusion, the two excised rings form a composite tissue that seals in a purse-string manner at both ends, similar to a single ring. In particular, the closure regions at both edges of the fused doublet contain aster-like (+1) defects in the organization of the actin fibers (Fig. 5).

**Figure 5.**
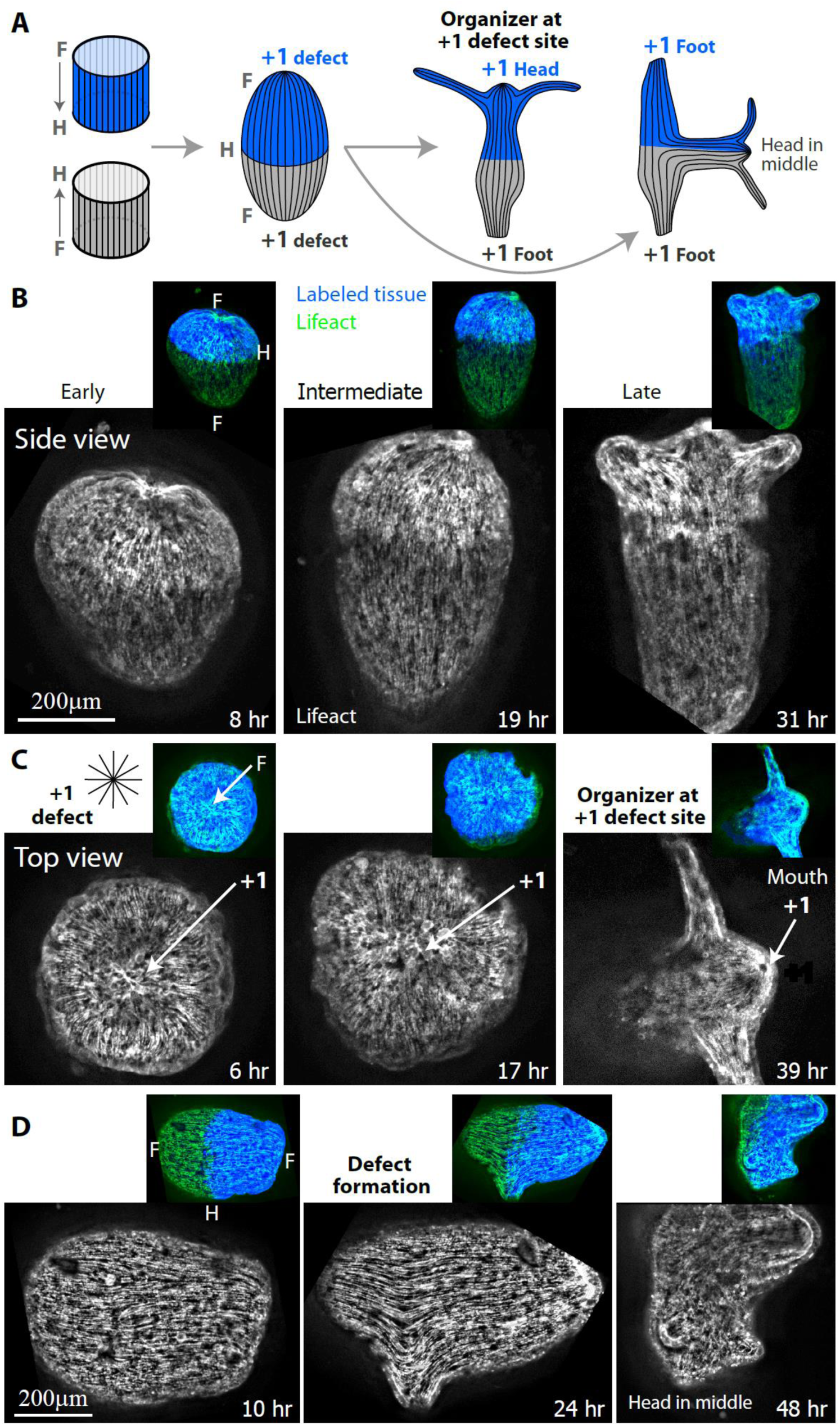
Actin fiber organization in regenerating H2H doublets. (A) Schematic illustration of the ectodermal actin fiber organization during regeneration of fused H2H ring doublets. Left: The actin fibers in the excised rings are arranged in parallel arrays, along the direction of the body axis of their parent animal. The two rings are positioned in a H2H configuration, and one of the rings is marked with a fluorescent tissue label (blue). Middle: Following fusion, the actin fibers from the two rings can join to form continuous fibers that span the length of the fused ring doublet. Two aster-like defects form at the top and bottom ends of the ring doublet. Right: In the case of polarity reversal (left; as in (B,C)), the fused ring doublet regenerates into an animal that has a normal morphology, with a new head forming at the aster-like (+1) defect site at one edge of the doublet, and a foot forming at the other end of the doublet. Alternatively, the H2H ring doublet can regenerate a new head in the middle, according to the original polarity of the excised rings (right; as in (D)). In this case, two feet form at the defects at the top and bottom ends of the fused tissue. (B,C) Spinning-disk confocal images from a time-lapse movie of a H2H doublet that underwent polarity reversal and regenerated into an animal with a head on an originally foot-facing side of one of the fused rings (Movie 8). Images are shown at frames providing a side view of the ring doublet where both rings are visible (B) or a top view of the tissue where only one ring is visible (C) at early (left), intermediate (middle) and late (right) time points during the regeneration process. The projected lifeact-GFP signal (see Methods) shows the organization of the ectodermal actin fibers. The defect at the edge is clearly visible from the top view in (C), and coincides with the formation site of the new organizer at the mouth of the regenerated animal. (D) Spinning-disk confocal images from a time-lapse movie of a H2H doublet that regenerated into an animal with a head in the middle and two feet at both edges (Movie 9). The projected lifeact-GFP signal depicting the organization of the ectodermal actin fibers is shown at an early (left), intermediate (middle) and late (right) time point during the regeneration process. An aster-like (+1) defect (surrounded by two -1/2 defects ^38^) forms *de novo* at the tip of a protrusion that appears at the middle of the regenerating doublet (middle) and develops into a head (right), similar to the process of bud formation ^45^. Insets in (B-D): Overlay depicting the lifeact-GFP signal (green) together with the fluorescent tissue label that marks one of the fused rings (blue; Texas Red-conjugated 3kD Dextran).

At the adhesion interface between the two rings, the excised tissues heal in a reasonably smooth manner. In particular, ectodermal actin fibers from the two rings, aligned along the common original axis of the parent animals, can form continuous cables that bridge across the interface between the two tissues in an ordered fashion with no apparent defects (Fig. 5A,B). In some cases, the adhesion interface is not fully organized into a continuous parallel array of fibers. Nevertheless, from the point of view of the overall organization of the actin fibers, the edges of the ring doublet with their associated aster-like defect are clearly distinct from the middle of the fused ring doublet that does not contain such defects.

The regeneration of fused ring doublets in the presence of frustration generates conditions in which polarity cannot be trivially maintained, and therefore provides a valuable opportunity to expose the contribution of the local structure to the establishment of a new organizer. We focus on regenerating H2H doublets where the formation of a head in the middle is suppressed (and hence the original polarity is not maintained), and find that the regenerated head emerges at the site of the preexisting aster-like defect at the doublet’s edge (Fig. 5A-C; Movie 8). As in a single ring, the aster-like defect at the edge forms rapidly as the parallel ectodermal fibers converge in the sealed wound, typically within ∼1-3 hours. However, in this case, the tissue region surrounding the defect was farthest from the head in the parent animal. The formation of a new head at an aster-like defect site at an originally aboral edge of a ring doublet is similarly observed during polarity reversal in the anti-oriented H2F configuration (Fig. S5). Importantly, in both of these cases in which head formation at the middle is suppressed, the alternative site at which a head appears is *not random* - head formation occurs precisely at the defect site at the edge of the composite tissue (Figs. 5C, S5). This does not intend to suggest that the defect itself is the “cause” or even the leading mechanism for the formation of the head organizer. Rather, we suggest that a combination of factors, including the emergence of bio-signals due to wound healing and the actin-fiber organization at the aster-like defect are dynamically coordinated to establish the new organizer. As such, under these circumstances, the presence of an aster-like defect can act reinforce head formation localizing the organizer precisely to the defect site.

The effect of structural factors on regeneration is also reflected in the suppression of head formation in the middle of ring doublets. While the oral side of individual rings invariably regenerates a head, in ring doublets where the oral side of one ring faces another tissue, head formation is often suppressed (Fig. 2F). This is most apparent in the oriented H2F configuration, in which nearly all samples regenerate into normal animals along their original polarity and suppress head formation in the middle (Fig. 2A-C). More surprising are the cases of polarity reversal in the anti-oriented H2F configuration, in which head regeneration occurs on the aboral side of the U ring, rather than the oral side of that ring at the middle. These observations suggest that the suppression of head formation in the middle of the tissue may be at least partly related to the absence of aster-like defects in the actin organization near the middle of fused ring doublets at the early stages of regeneration; the tissue at the middle interface which lacks a defect provides a less favorable substrate for head formation. Interestingly, a head can also emerge in the middle of a fused ring doublet in a region that initially lacks defects (Fig. 5A,D; Movie 9). This is most common in H2H ring doublets where the memory of polarity introduces a strong bias for head formation in the middle (Fig. 1). In cases where this bias results in middle head formation, the emergence of a head is accompanied by reorganization of the actin fibers that involves the *de-novo* formation of defects in the actin fibers at later stages of the regeneration (Fig. 5D). The two defects at the edges of the composite tissue in this case will become feet, as expected from the original polarity of the tissues. The emergence of a head in the middle likely involves biochemical signaling that drives the mechanical processes leading to tissue deformations and reorganization of the cytoskeleton, similar to what occurs during bud formation ^45^. As in emerging buds, the initiation of the structural protrusion in head formation at the middle coincides with the reorganization of the cytoskeleton and the formation of defects at the protrusion site (Fig. 5D).

## Discussion

While the inductive properties of an existing head organizer are evident, the way a new organizer emerges during regeneration of excised *Hydra* tissues that initially lack a functional organizer is still elusive. Given that the entire gastric tissue in *Hydra* has the potential to form a new organizer, what directs head formation to a particular site? How is the information instilled in the excised tissue linked to the establishment of a polar body axis in the regenerated animal? To probe these questions, we studied the establishment of polarity in regenerating ring doublet tissues assembled in various frustrating configurations. The regeneration under these conflicting circumstances exposes the dynamic nature of polarity determination, and allowed us to explore the interplay between the different processes involved.

The dominant view of polarity patterning in *Hydra* regeneration is based on memory; the inheritance of biochemical gradients from the parent animal ^13, 14^. According to this view, after removal of the head and foot regions, a regenerating tissue establishes a new organizer region by a local self-enhancing reaction that is coupled with a long-range inhibitory effect. The initiation of local activation process depends primarily on the preexisting activation properties of the tissue which are graded as a function of distance from the original organizer ^6, 14^. In addition to these preexisting activation properties, the injury generated by the excision plays a central role in head formation ^33^, inducing a strong response at the wound site ^10, 26, 27, 34, 35^.

Interestingly, recent reports show that the injury response is generic, with similar patterns of signaling, gene expression, and apoptosis appearing at oral and aboral-facing wounds initially, ^10, 27, 35^. Nevertheless, an excised tissue ring or bisected animal robustly preserves their original polarity despite (Fig. S1). If the injury response is symmetric, what generates the asymmetry that robustly leads to regeneration of different structures at the oral and aboral end? What relays the positional information from the parent animal? Despite extensive research, the answers to these questions remain unknown, emphasizing that the link between the tissue memory based on activation gradients in the parent animal, the injury response, and the subsequent patterning process, is still far from understood.

The ring doublet experiments provide a well-controlled platform to study these questions. We observe a striking contrast between configurations with an initial bias where ring doublets reproducibly regenerate according to the inherited polarity memory on the one hand, and frustrated configurations where we observe variable morphological outcomes including polarity reversal on the other hand. The reproducibility of the regeneration patterns in ring doublets with an initial bias is evident in the oriented H2F configuration, where nearly all tissues regenerate normally and maintain their original polarity (Fig. 2B top). Interestingly, this is true despite the inherent variation between parent animals, indicating that the graded positional information in tissues excised from different animals scales with the size of the animal. Similarly, in the asymmetric F2F ring doublets, all samples regenerate a head at the oral-facing wound that was closer to the head organizer in the parent animals (Fig. 3). Thus, the memory of tissue position and polarity is effectively retained even in thin tissue rings (∼10-20% of the body column), after the removal of the head organizer and the foot signaling center, and these initial biases are sufficient for directing the morphological outcome during regeneration in a highly reproducible manner. Nevertheless, in frustrated configurations (e.g. in the H2H or anti-oriented H2F configurations), the ring doublets exhibit variable morphological outcomes (Fig. 1, Fig. 2B bottom), demonstrating that polarity memory is labile and that the site of head formation is not preset but rather emerges dynamically in the regenerating tissue. The strong dependence of the morphological outcome at an oral or aboral wound on the entire tissue (as evident from the sensitivity to the doublet configuration) is a signature of long-range information transfer across the tissue. Moreover, the reproducibility of the morphological outcomes in biased doublet configurations further suggests that the observed variability under frustrating conditions is due to a dynamic competition between sites, rather than random stochasticity or variations associated with the experimental manipulations involved in sample preparation.

We emphasize the difference between the ring doublet experiments and the more-commonly used approach in which regeneration is studied following bisection (i.e. where the regenerating tissue retains either the head or the foot of the parent animal). In bisected animals, the site of head or foot formation is predictable and the long-range influence of the preexisting head organizer or foot signaling center likely dominates the patterning process ^10^. In contrast, the location of a new head organizer in regenerating ring doublets under frustration is an emerging property of the tissue that is not predetermined in an obvious way at the onset of the process. As such, the ring doublets facilitate the investigation of subtle differences in the dynamic processes leading to the establishment of a polar body axis that may be masked in bisected animals.

The effective competition between different potential sites for head formation implies sensitivity to initial conditions; slight variations at early times (e.g. due to the history of the tissue or its structure) can be amplified to direct head formation to a particular region and thus steer the polarity of the regenerated animal. In other words, head formation reflects a complex dynamic competition between different tissue regions that have the potential to support the development of a head organizer, rather than a preset patterning process that specifies a particular site. Importantly, this dynamic view emphasizes the plasticity of the regeneration process, whereby a multi-faceted integration of multiple factors, rather than a direct causal relation, determines the polarity of the regenerated animal. Such a dynamic view of the regeneration process can rationalize the cohort of our experimental observations in the different configurations studied here. This view is also compatible with recent reports showing that regeneration is initiated via a strong position-independent injury response that cannot predict the eventual regeneration trajectory ^10^. The diversification in the patterning process that eventually leads to the formation of different structures is apparent only much later in the process and involves complex interactions between multiple signaling pathways and possibly other factors.

We coupled our investigation of the plasticity of body axis polarity in regenerating ring doublets under frustration with a detailed examination of their cytoskeletal organization. This was motivated by our hypothesis that tissue mechanics, and in particular the actomyosin force generating system, could be an integral part of the feedback loops involved in polarity determination and stabilization ^39^. We show that head formation occurs preferably (but not exclusively) at the edge of fused ring doublets, which after closure invariably contains an aster-like defect in the alignment of the supra-cellular ectodermal actin fibers (Fig. 5). Strikingly, this is true even when polarity is reversed, i.e. when a head forms at the originally aboral-facing edge of one of the fused tissue rings (Fig. 3E). Furthermore, we observe suppression of head formation at the interface between the two rings that initially lacks an aster-like defect, even though it contains an oral-facing wound (Fig. 3F).

The biochemical and mechanical processes involved in polarity determination and head formation are intertwined, with extensive crosstalk and mutual feedback which eventually lead to the formation of a functional head (possibly also with the involvement of electrical processes, shown to play a role in wound healing and regeneration ^47^). The new head will invariably contain an organizer region which acts as a signaling center and has a characteristic cytoskeletal arrangement with an aster-like defect in the ectodermal actin fibers. Is this stereotypical cytoskeletal organization a downstream corollary of the signaling processes associated with the establishment of a new organizer? Or could the organization of the cytoskeletal be an integral part of the patterning processes that eventually lead to the establishment of a stable organizer?

The Wnt signaling pathway is known to be a key component of the head organizer in *Hydra* that can induce new head formation ^3, 6, 8, 30^. Indeed, global activation of the Wnt pathway leads to the formation of multiple heads ^30^. Moreover, local enhancement of Wnt signaling can direct organizer formation and override structural factors ^23^. Recent work has also shown that Wnt signaling is activated at wound sites ^10, 27, 35^. However, the initial activation patterns of Wnt signaling during injury appear generic, irrespective of the morphological outcome. In particular, aboral wounds in bisected animals which develop into feet show similar early Wnt3 activation as oral wounds which will be the site of the new head organizer. In that respect, it is also worth noting that overexpression of Wnt3 might lead to different dynamics than the “natural” ones following the spontaneous self-organization within a regenerating tissue.

We suggest that the specialized biochemical, mechanical and structural properties at the head organizer site in regenerating tissue pieces can arise via different scenarios, which can be observed under different circumstances. In one scenario, in the absence of a preexisting aster-like defect, autocatalytic signaling events initiate head formation by creating the necessary cytoskeletal organization and mechanical conditions. This will lead to *de novo* defect formation as seen in middle-head formation in the H2H configuration (Fig. 5D). This scenario also appears to occur in bud formation ^45^ or regeneration from tissue pieces with an initial bias in Wnt concentrations ^23^. In all these cases, the hierarchy of events is clear; the aster-like defect is induced by local cues generated by signaling processes which direct head formation.

In an alternative scenario, an aster-like defect emerges in the tissue early on, within timescales overlapping with the effects of injury response and other signals. This occurs e.g., at the edge of a regenerating ring doublet, where an aster-like defect in the ectodermal myonemes forms within 1-3 hours at the wound site at the edge (Fig. 5). Given the extensive crosstalk between the Wnt signaling pathway and mechanical processes ^48^, the formation of a new organizer will likely depend on mutual feedback and mechanical cues. In particular, the local mechanical environment at the defect site can provide mechanical cues that help localize the organizer to that particular region. In this sense, the defect acts can actively reinforce organizer formation. This scenario seems relevant, for example, at defects identified early during polarity reversal in regenerating ring doublets (Figs. 5B,C, S5, Movie 8) or in regenerating tissue fragments ^38^. This is also compatible with our recent experiments on tissue strips ^49^, showing the emergence of an aster-like defect following the folding of the tissue segment into a hollow spheroid and the emergence of an organizer at the same site without the existence of an explicit preexisting signaling source.

We emphasize that in this alternative scenario, given the extensive crosstalk between the biochemical and mechanical processes, it is difficult to describe organizer formation as a simple causal chain of events. In particular, neither the presence of an aster-like defect nor the early activation of the Wnt-signaling pathway are reliable pre-indicators for head formation. This is evident, e.g. in the aboral end of a ring or bisected animal which contains a defect (Fig. 4) and also exhibits local activation of the Wnt signaling pathway ^10, 27, 35^, but nonetheless develops into a foot. These observations suggest that it may not be possible to assign a well-defined single causal driver for head formation in these cases. Rather, the interplay between signaling processes and mechanical feedback, that also involves long-range information transfer reflecting the context-dependent global state of the tissue, eventually leads to the formation a new organizer in a dynamic self-organization process which directs the regeneration of a functional animal.

The observed bias favoring organizer formation at aster-like defect sites in regenerating ring doublets, is well-aligned with recent results by us and others, that identified the aster-like defects (i.e.+1 nematic topological defects) as organization centers for head formation in regenerating *Hydra* tissue pieces ^38, 49^ and mouth formation in regenerating jellyfish ^50^. The inherent structure of these defects as focal points of multiple contractile actin fibers implies that these sites are mechanically unique ^38^. Moreover, once formed, these defects were shown to remain stationary relative to the underlying cells ^38^. As such, the cells near the core of the defect experience a sustained distinct mechanical environment that can locally activate signaling pathways, effectively turning this site into an attractor for head formation. Indeed, observations in regenerating jellyfish showed that local activation of the Wnt pathway occurred specifically at aster-like defects sites ^50^, directly demonstrating the coupling between cytoskeletal organization, mechanics and relevant signaling pathways. Thus, while a defect is neither a prerequisite nor necessarily associated with head formation, its presence can nonetheless have a profound impact on head formation through its intimate coupling to signaling dynamics.

A central feature of the establishment of the head organizer is the autocatalytic nature of the local activation. While the Wnt pathway has been shown to be self-activated ^28^, we propose that the mechanical environment in the vicinity of an aster-like defect can also provide sustained local activation promoting the formation and stabilization of the head organizer. Earlier work has shown that all cells in *Hydra* turn over, with cell removal occurring at tip of the hypostome, the tip of the tentacles and the peduncle at the foot ^51^. We can now identify all these sites of cell extrusion in the mature *Hydra* as sites containing aster-like (+1) defects in the actin organization ^38^, in analogy with recent observations *in vitro* where cell extrusion in epithelial sheets was localized at defect sites ^52^. Due to the stability of the +1 defects and the unique mechanical environment at defect sites ^38^, the presence of a defect can be part of the feedback loops that define the head organizer and contribute to its stabilization in the presence of continuous tissue renewal ^6^ and through it to the integrity of the *Hydra* body plan over time.

The large body of work on *Hydra* polarity starting from Ethel Browne more than a 100 years ago and continued by many others over the years, has led to many insights. Yet many basic questions remain open: what are the factors responsible for positional memory in the tissue? How are these factor related to the injury response at the wound site? What is responsible for the long range information transfer across the tissue? How are local effects integrated with tissue scale processes to direct the formation a new head organizer? While we still cannot answer all these questions, our work puts forward a dynamic view of polarity determination. Rather than seeking a specific molecular precursor or a well-defined causal trajectory for head regeneration, we claim that the formation of a new head organizer arises from a multi-faceted integration of diverse contributions that can proceed along different developmental paths.

Furthermore, while the current views of polarity emphasize diffusible biochemical signals as the main carriers of polarity information, we suggest that additional factors such as mechanical processes can contribute to the establishment of a polar body axis ^38, 39, 49^. More broadly, we believe that such an integrated framework that takes into account the symbiotic interactions between the different mechanical and biochemical factors will be essential for deciphering the mechanisms responsible for polarity determination in animal morphogenesis.

## Methods

### *Hydra* Strains and Culture

Experiments are performed with a transgenic strain of *Hydra* Vulgaris (AEP) expressing lifeact-GFP in the ectoderm ^53^ (generously provided by Bert Hobmayer, University of Innsbruck). Animals are cultivated in *Hydra* medium (HM; 1 mM NaHCO3, 1 mM CaCl2, 0.1 mM MgCl2, 0.1 mM KCl, and 1 mM Tris-HCl, pH 7.7) at 18° C. The animals are fed three times a week with live *Artemia nauplii* and rinsed after 4 hr. Experiments are initiated ∼24 hr after feeding.

### Fluorescent Tissue Labeling

In order to distinctly label tissue segments originating from different parent *Hydra* we fluorescently label the animals (prior to excising rings) by electroporating fluorescent cell volume dyes. Electroporation of the dye into live *Hydra* is performed using a homemade electroporation setup ^38^. The electroporation chamber consists of a small Teflon well, with 2 perpendicular Platinum electrodes, spaced 5 mm apart, on both sides of the well. Four *Hydra* are placed in the chamber in 20μl of *Hydra* medium supplemented with 2mM of dye (Alexa Fluor 647-conjugated 10kD Dextran or Texas Red-conjugated 3kD Dextran, both from Invitrogen). A 150 Volts electric pulse is applied for 35ms. The animals are then rinsed in *Hydra* medium and allowed to recover for ∼12 hours prior to tissue excision. Unless stated otherwise, experiments involving fusion of two *Hydra* tissue rings are performed using two separate donor animals, each labeled with a different colored dye.

To label the oral region of a single excised ring we use laser photoactivation of a caged dye (Abberior CAGE 552 NHS ester) that is electroporated uniformly into mature *Hydra* and subsequently activated in the desired region, as previously described ^38^. The electroporation chamber consists of 2 perpendicular Platinum electrodes, spaced 2.5 mm apart. A single *Hydra* is placed in the chamber in 10μl of HM supplemented with 4mM of the caged dye. A 75 Volts electric pulse is applied for 35ms. The animal is then washed in HM and allowed to recover for several hours to 1 day prior to tissue excision. Following excision, the specific region of interest is activated by a UV laser in a LSM 710 laser scanning confocal microscope (Zeiss), using a 20× air objective (NA=0.8). Photoactivation of the caged dye is done using a 10 mW 405nm laser at 100 %. Subsequent imaging of the lifeact-GFP signal and the uncaged cytosolic label is done by spinning-disk confocal microscopy as described below. The uncaged dye remains within the cells of the photoactivated region and does not get advected or diffuses away within our experimental time window ^38^.

### Sample Preparation

#### Extracting rings with known polarity with respect to parent animal

In order to investigate the influence of the original body axis polarity on the polarity of the regenerated samples, it is necessary to excise tissue rings in a manner which preserves knowledge of their original polarity. For this purpose, we use a platinum wire (diameter: 75 μm, length: ∼1 cm), bent at one end to provide a directional marker. Next, we cut off one side of the animal (head or foot) and insert the straight end of the wire into the body cavity (along its axis). An additional transverse cut removes the other side of the body (foot or head, respectively), and leaves us with a ring threaded on the wire with known polarity. To study polarity preservation in individual rings (Fig. S1), the samples are kept on the wire and transferred to a 96 well plate for imaging. The rings are excised using a scalpel equipped with a #15 blade.

#### Preparation of fused ring doublets

To generate samples with frustrated initial conditions, we fuse two tissue rings to each other. We use different configurations with regards to the original polarity (H2H, H2F, F2F) and the relative body positions of the two rings (UU, LL, LU, UL). We define the relative body position in the parent animal by dividing the *Hydra* body into four parts: Head, Upper (U), Lower (L) and Foot. In all experiments we use only the U and L ring sections, positioned above and below the estimated middle of the animal, respectively. The length of each tissue ring (along the animal axis) ranges from about 1/8 to 1/4 of the full animal’s length.

A fused ring doublet is generated by combining two rings, excised from two separate animals that are differentially labeled by a fluorescent dye (see above). The first ring is excised from the desired position (U or L) in one *Hydra* and threaded on a wire in the desired orientation as described above. The excised ring is then transferred, while maintaining its orientation and threaded onto a vertical 75 μm-diameter platinum wire which sticks out of a petri dish that is half-filled with 2% agarose (prepared in HM) and layered with additional HM from above. The procedure is repeated with the second ring (taken from another animal) which is placed on top of the first ring (so that both rings are immersed in HM). Although close contact for a few minutes is sufficient for establishing an initial connection between the two ring tissues, significantly longer time is needed to ensure stable fusion. An additional horizontal wire is placed on top of the two rings to hold them together. We did not observe differences in the regeneration outcome as a function of the overall orientation of the ring doublet on the vertical wire (i.e. which ring was placed on the bottom or the top of the vertical wire, as long as the overall doublet configuration was maintained).

The tissue rings are left to adhere to each other for 6-8 hours on the vertical wire. Subsequently, the wires are removed, and samples consisting of two rings that have fused together form composite ring doublets. Typically, 12-24 samples are generated in one experiment. The samples are transferred into a 25-well agarose setup placed in a 50 mm glass-bottom petri dish (Fluorodish) for imaging. The wells are produced using a homemade Teflon 25-pin comb (1.5 mm diameter pins) with 2% agarose in HM. Each sample is placed in a well which is filled with HM. The tissue originating from each ring can be identified by its distinct fluorescent label (from the parent animal labeled with a different dye). This allows us to track the dynamics of the tissues originating from each of the two rings in the regenerating fused ring doublet. Using this protocol ∼90% of the samples fuse successfully.

#### Fused ring-doublets formed with a physical barrier in the middle

To introduce a physical barrier at the interface between the two rings forming the doublet, we insert a horizontal platinum wire (diameter: 75 μm) as an obstacle between the two rings. The procedure for preparing these samples is similar to the procedure for preparing the fused ring doublets described above, except for the insertion of the horizontal wire at the fusion interface between the two rings. The horizontal wire is placed on the bottom ring, along the interface between the rings, before placing the second ring on it. The two rings adhere to each other around the horizontal wire (on both sides), while remaining threaded on the vertical wire that holds them in place. After ∼6-8 hours the wires are removed after checking carefully that the horizontal wire was indeed pinned between the two rings (by checking the sample can be lifted using this wire). The success rate of fusion with a barrier in this geometry is ∼50%.

#### Microscopy

Time-lapse epifluorescence and bright field movies of regenerating *Hydra* are recorded on a Zeiss Axio-Observer microscope with a 5x air objective (NA= 0.25), at room temperature. Images are acquired on a charge-coupled device (CCD) camera (CoolSnap, Photometrix), and illuminated with an XCite 120Q lamp (Excelitas Technologies). Time lapse imaging begins after the ring doublets are removed from the wire (∼6-8 hours after excision) and continues for 3 days at a time interval of 10-15 minutes. We use 4 channels: bright field, Lifeact-GFP, Dextran Texas Red, Dextran Alexa Fluor 647.

Time-lapse spinning-disk confocal movies are acquired on a spinning-disk confocal microscope (Intelligent Imaging Innovations) running Slidebook software. Imaging is done using laser excitation at 488 nm (lifeact-GFP), 561 nm (Texas Red) and 640 nm (Alexa-fluor 647) and appropriate emission filters at room temperature and acquired with an EM-CCD (QuantEM; Photometrix). In some cases, to reduce tissue movements and rotations, the sample is embedded in a soft gel (0.5% low melting point agarose (Sigma) prepared in HM). Time lapse movies of regenerating *Hydra* are taken using a 10× air objective (NA=0.5). Due to light scattering from the tissue, we cannot image through the entire sample so our observations are limited to the side of the tissue that is facing the objective. Spinning-disk z-stacks are acquired with a 3-5 μm interval between slices for a total of 120 μm.

Final images of the regenerated *Hydra* are taken after 3-5 days, after relaxing the animals in 2% urethane in HM for 1 minute and sandwiching them between two glass coverslips with a 200 μm spacer between them. The samples are imaged from both sides (by flipping the sample).

#### Analysis

The outcome morphology of regenerated ring doublet samples is determined manually based on inspection of the time-lapse movie of the regeneration process and the final images. Regeneration is defined as the formation of a head structure with tentacles within 3 days. The regenerated samples develop a variety of different morphological structures (Fig. S2). For each doublet configuration, the outcome morphologies of the different samples are divided into several categories (as indicated schematically in the respective bar plots) based on the appearance of morphological structures head (with tentacles)/foot and their position with respect to the fused ring tissues (as determined from the distribution of the labeled tissues originating from each ring). This allows us to determine if structures such as a regenerated head originate from the interface between the two rings (and are hence labeled with both colors), or from either side of the ring doublet. The tissue originating from the head or foot facing sides of the excised rings can be identified based on the distribution of the tissue labels and the knowledge of the doublet’s configuration. Polarity reversal refers to situations in which a regenerated head forms on the originally foot-facing side of one of the rings. Note that this tissue region would invariably be the site of foot formation if the ring was allowed to regenerate on its own (Fig. 4).

To obtain an estimate of the sizes of the two rings in a fused ring doublet, we measured the projected area of the tissue originating from each ring in the fused spheroid. To obtain a reliable estimate of the relative sizes of the tissue originating from the two rings, we choose a frame from the time-lapse movie in which the interface between the two rings is roughly perpendicular to the imaging plane. Furthermore, since the regeneration process involves a series of osmotic inflations and abrupt shrinkages, we choose to measure the projected area shortly after a shrinking event to obtain a more time-independent estimate of the projected areas.

The organization of the supra-cellular actin fibers in the ectoderm is followed by analyzing time lapse spinning disk confocal movies of the lifeact-GFP signal in regenerating samples, in addition to the labels marking the tissue originating from the different rings. The projected images of the supra-cellular actin fibers (myonemes) in the basal layer of the ectoderm are extracted from the 3D z-stacks at each time point as described in ^38^.

## Acknowledgments

We thank Gidi Ben Yoseph for superb technical assistance. We thank Prof. Bert Hobmayer for generously providing transgenic *Hydra* expressing lifeact-GFP. We thank Omri Wurtzel and Tom Schultheiss for valuable discussions. We thank Omri Wurtzel and Niv Ierushalmi for comments on the manuscript. This work was supported by a grant from the European Research Council (ERC-2018-COG grant 819174) to K.K., a grant from the Israel Science Foundation (grant No. 228/17) to E.B., and a Miriam and Aaron Gutwirth Memorial Fellowship to Y.M.S.

## Supplementary Information

### Supplementary movies

**Movie 1. H2H doublet that regenerated into an animal with a head in the middle.** Time-lapse movie of a H2H doublet that underwent polarity reversal, regenerating into an animal with a head in the middle and two feet at the edges. Combined epifluorescence (Left; green-AlexaFluor 647-conjugated 10kD dextran, magenta-Texas Red-conjugated 3kD dextran) and bright-field (Right) images are shown. The elapsed time from excision is displayed (hrs:min), and the scale bar is 200 µm.

**Movie 2. H2H doublet that regenerated a head on its edge.** Time-lapse movie of a H2H doublet that underwent polarity reversal, regenerating into an animal with a normal morphology, with a head that developed from an originally foot-facing side of one of the excised rings. Combined epifluorescence (Left; green-AlexaFluor 647-conjugated 10kD dextran, magenta-Texas Red-conjugated 3kD dextran) and bright-field (Right) images are shown. The elapsed time from excision is displayed (hrs:min), and the scale bar is 200 µm.

**Movie 3. H2F doublet in the oriented configuration that regenerated along its original polarity.** Time-lapse movie of an oriented H2F doublet that regenerated into a normal animal along the original body axis orientation. Combined epifluorescence (Left; green-AlexaFluor 647-conjugated 10kD dextran, magenta-Texas Red-conjugated 3kD dextran) and bright-field (Right) images are shown. The elapsed time from excision is displayed (hrs:min), and the scale bar is 200 µm.

**Movie 4. H2F doublet in the anti-oriented configuration that reversed polarity.** Time-lapse movie of an anti-oriented H2F doublet that regenerated into an animal whose body axis is oriented in the opposite direction to its original polarity, with a head forming on the original foot-facing edge. Combined epifluorescence (Left; green-AlexaFluor 647-conjugated 10kD dextran, magenta-Texas Red-conjugated 3kD dextran) and bright-field (Right) images are shown. The elapsed time from excision is displayed (hrs:min), and the scale bar is 200 µm.

**Movie 5. F2F doublet that regenerated into a normal morphology with a single head.** Time-lapse movie of a F2F ring doublet that regenerated into an animal with a single head on the U side. Combined epifluorescence (Left; green-Texas Red-conjugated 3kD dextran, magenta-AlexaFluor 647-conjugated 10kD dextran) and bright-field (Right) images are shown. The elapsed time from excision is displayed (hrs:min), and the scale bar is 200 µm.

**Movie 6. F2F doublet that regenerated two heads.** Time-lapse movie of a F2F ring doublet that regenerated into an animal with two heads on both sides. Combined epifluorescence (Left; green-Texas Red-conjugated 3kD dextran, magenta-AlexaFluor 647-conjugated 10kD dextran) and bright-field (Right) images are shown. The elapsed time from excision is displayed (hrs:min), and the scale bar is 200 µm.

**Movie 7. Actin organization in a single regenerating ring.** Time-lapse, spinning-disk confocal movie depicting the actin organization in a regenerating ring. The oral side of the ring was labeled with a fluorescent tissue label (by locally uncaging Abberior CAGE 552) to mark the original polarity of the tissue, which is preserved in the regeneration process. Right: projected lifeact-GFP signal showing the organization of the ectodermal actin fibers. Left: overlay of the lifeact-GFP signal (green) with the fluorescent tissue label (blue). The elapsed time from excision is displayed in hours (hrs:min), and the scale bar is 100 µm.

**Movie 8. Actin organization in H2H doublet that underwent polarity reversal.** Time-lapse, spinning-disk confocal movie depicting the actin organization in a H2H doublet that underwent polarity reversal and regenerated into an animal with a head on an originally foot-facing side of the labeled ring. The aster-like defect at the labeled edge is clearly visible from the beginning of the movie, and coincides with the site of head formation in the regenerated animal. Right: projected lifeact-GFP signal showing the organization of the ectodermal actin fibers in the ring doublet. Left: overlay of the lifeact-GFP signal (green) with the fluorescent tissue label marking one of the rings (blue; Texas Red-conjugated 3kD dextran). The elapsed time from excision is displayed in hours (hrs:min), and the scale bar is 100 µm.

**Movie 9. Actin organization in H2H doublet that regenerated a head in the middle.** Time-lapse, spinning-disk confocal movie of a H2H doublet that regenerated into an animal with a head in the middle and two feet at both edges. A +1 defect (surrounded by two -1/2 defects) forms *de novo* at the tip of a protrusion that appears at the middle of the regenerating doublet and develops into a middle head. Right: images of the projected lifeact-GFP signal showing the organization of the ectodermal actin fibers in the ring doublet. Left: overlay of the lifeact-GFP signal (green) with the fluorescent tissue label marking one of the rings (blue; Texas Red-conjugated 3kD dextran). The elapsed time from excision is displayed in hours (hrs:min), and the scale bar is 100 µm.

### Supplemental Figures

**Figure S1.**
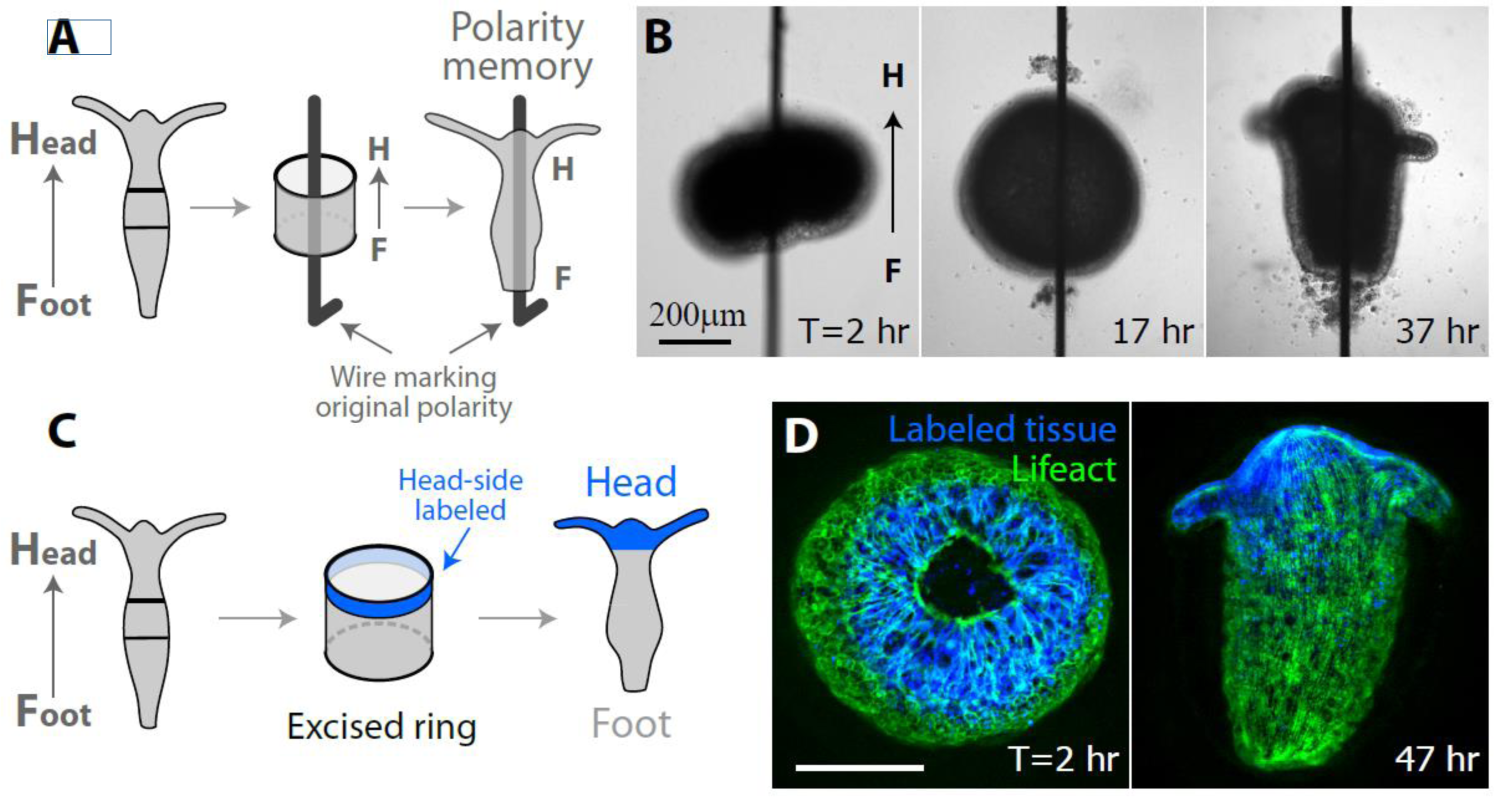
Memory of polarity in regenerating single tissue rings. (A,B) Memory of polarity in single tissue rings regenerating on a wire. (A) Schematic illustration of the regeneration of an excised tissue ring on a wire ^37^. A ring is excised from the gastric region of a mature *Hydra*. The ring is threaded on a wire that is bent at one end to mark the original polarity of the tissue. The majority of excised rings observed were able to regenerate on the wire (61 out of 92 samples regenerated, 12 escaped from the wire, 8 disintegrated, and 11 did not regenerate). 60 out of the 61 regenerated samples had a normal morphology that maintained the original polarity of the excised ring (1/61 samples regenerated into an abnormal morphology). (B) Images from a time lapse movie of an excised ring that regenerated on a wire. The original polarity of the excised ring is maintained in the regenerated animal. (C,D) Memory of polarity in a regenerating tissue ring that is labeled at its oral side. (C) Schematic illustration of the regeneration of a tissue ring excised from the gastric region of a mature animal. The excised ring regenerates into an animal with a new head at the labeled region and a new foot at the other end, thus maintaining its original polarity. (D) Spinning-disk confocal images from a time-lapse movie of an excised ring that is labeled at its oral end (Movie 7). The labeling is done by locally uncaging an electroporated caged-dye (Abberior CAGE 552; see Methods) ^38^. Images show an overlay of the projected lifeact-GFP signal (green) and the fluorescent tissue label (blue) that marks the tissue originating from the oral edge of the tissue ring shortly after excision (left) and in the regenerated animal (right).

**Figure S2.**
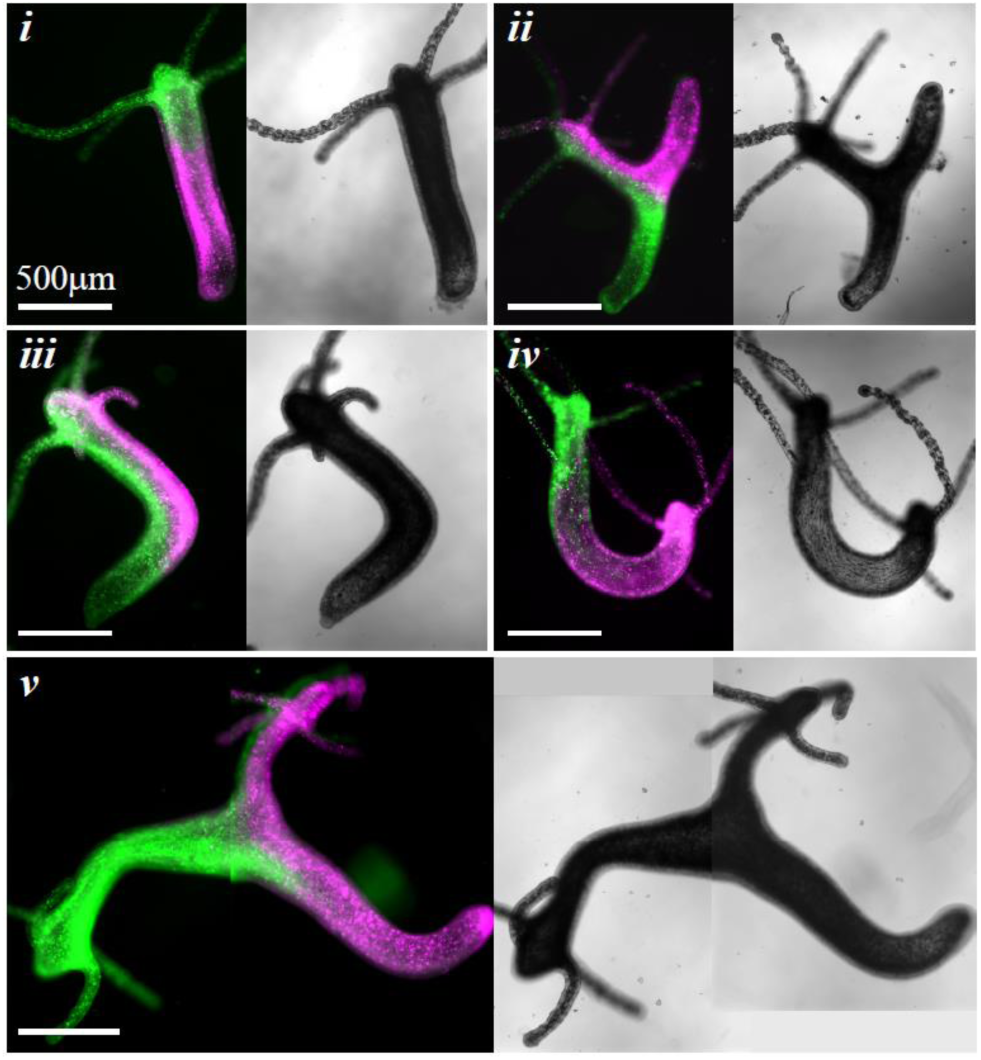
Different types of morphological outcomes in regenerating ring doublets. Combined epifluorescence (Left; green-AlexaFluor 647-conjugated 10kD dextran, magenta-Texas Red-conjugated 3kD dextran) and bright-field (Right) images of different outcome morphologies observed in the regeneration of ring doublets in the various configurations: (i) Normal (ii) Head in the middle with two feet (iii) Head originating from the middle of the doublet with normal morphology (iv) Two heads at opposite sides and (v) Heads in the middle and on the side.

**Figure S3.**
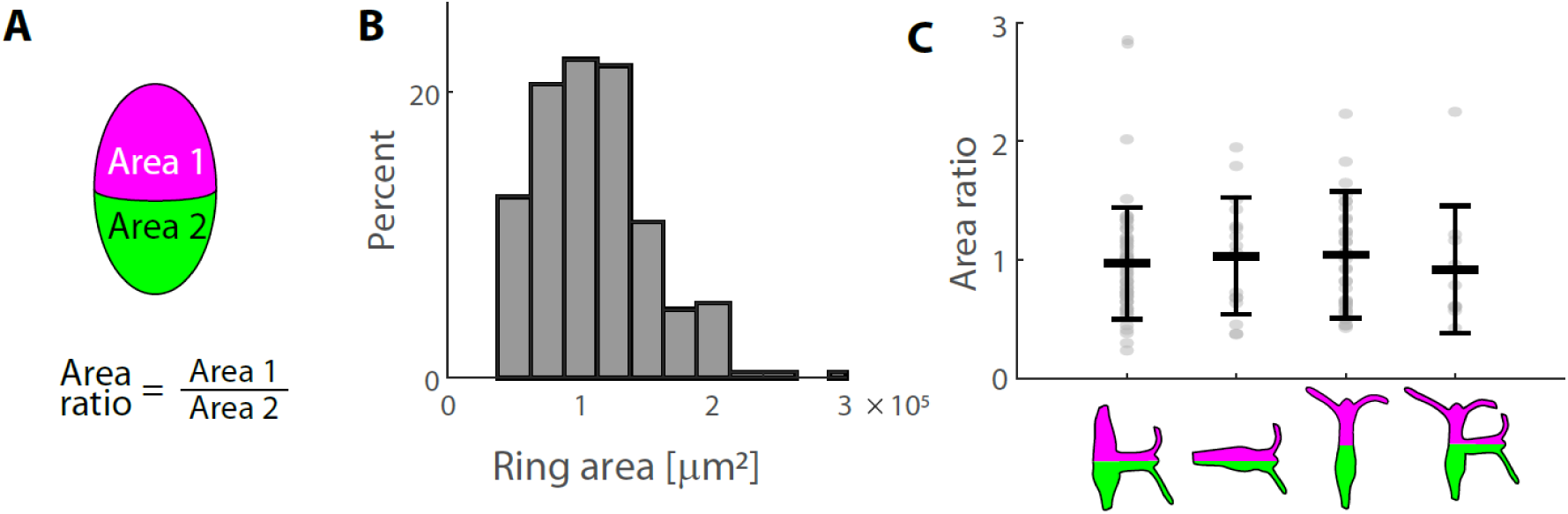
The relative sizes of fused tissue rings in H2H doublets. The size of the tissue originating from the two rings was characterized by the projected area within the fused spheroid (N=114 H2H doublets; see Methods). (A) Schematic illustration showing the projected areas, and the area ratio which provides a measure of the relative sizes of the two rings. (B) Bar plot showing the size distribution of projected ring areas (N=228 rings). (C) A graph depicting the area ratio of regenerating H2H ring doublets as a function of the possible morphological outcomes. For each type of outcome, the mean area ratio and standard deviation are shown (black), as well as a scatter plot of the area ratio values for each ring doublet (gray dots).

**Figure S4.**
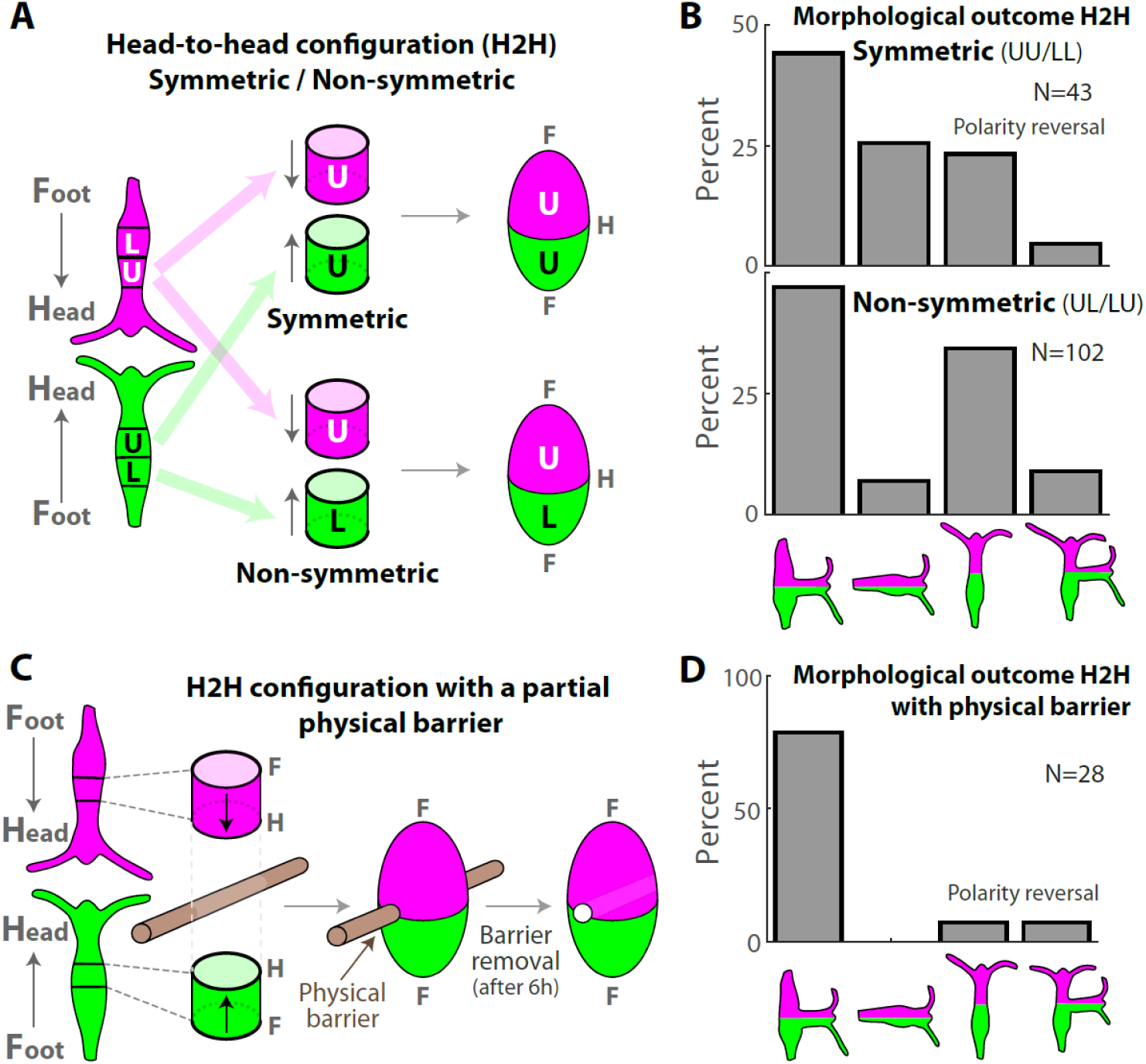
Detailed analysis of the regeneration of H2H doublets. (A) Schematic illustration of the regeneration experiments with fused ring doublets in the H2H configuration. Rings are excised from above (U) or below (L) the approximated midpoint of the two parent animals that are differentially labeled (green/magenta). The rings are fused so that their originally head-facing sides adhere to each other. Samples are made from two rings taken from the same position along the body axis of the parent animals (top; symmetric UU or LL) or from different positions (bottom; non-symmetric UL or LU). (B) Bar plot depicting the outcome morphologies of fused H2H ring doublets generated in a symmetric (UU or LL; top) or non-symmetric (UL or LU; bottom) manner. (C) Schematic illustration of the formation of a H2H doublet with a physical barrier (a 75 μm-diameter wire) placed at the adhesion site. The barrier is removed after ∼6 hours and the regeneration proceeds as in (A). (D) Bar plot depicting the outcome morphologies of H2H doublets formed with this physical barrier. The probability for polarity reversal with head formation at an originally foot-facing side of one of the excised rings (right bars) is reduced by the presence of the barrier (compare to (B)).

**Figure S5.**
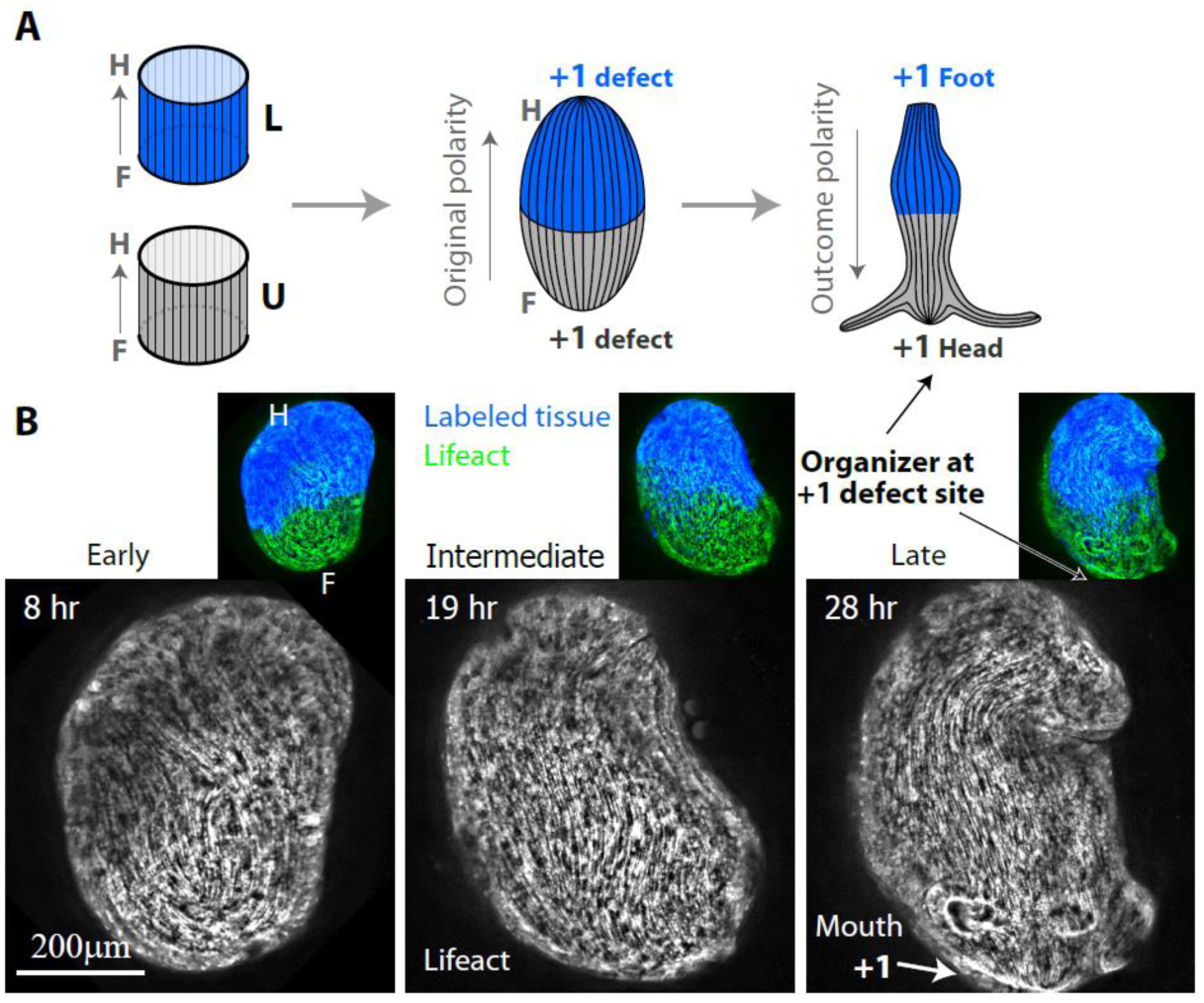
Actin fiber organization in a regenerating H2F ring doublet that undergoes polarity reversal. (A) Schematic illustration of the ectodermal actin fiber organization during regeneration of an anti-oriented H2F ring doublet that undergoes polarity reversal. Left: The actin fibers in the excised rings are arranged in parallel arrays, along the direction of the body axis of their parent animal. The two rings are positioned in an anti-oriented H2F configuration, and one of the rings is marked with a fluorescent tissue label (blue). Middle: Following fusion, the actin fibers from the two rings can join to form continuous fibers that span the length of the fused ring doublet. Two aster-like defects form at the top and bottom ends of the ring doublet. Right: In the case of polarity reversal, the fused ring doublet regenerates into an animal that has a normal morphology, with a new head forming at the aster-like (+1) defect site at the bottom (originally foot-facing) edge of the doublet, and a foot forming at the other (originally head-facing) end. (B) Spinning-disk confocal images from a time-lapse movie of an anti-oriented H2F doublet that underwent polarity reversal. Images are shown at an early (left), intermediate (middle) and late (right) time points during the regeneration process. The projected lifeact-GFP signal (see Methods) shows the organization of the ectodermal actin fibers. The +1 defect site at the bottom edge of the sealed doublet coincides with the formation site of the new organizer at the mouth of the regenerated animal. Insets: Overlay depicting the lifeact-GFP signal (green) together with the fluorescent tissue label that marks one of the fused rings (blue; Texas Red-conjugated 3kD Dextran).

